# Identifying small molecule probes of ENTPD5 through high throughput screening

**DOI:** 10.1101/503938

**Authors:** Matthew A. Durst, Kiira Ratia, Arnon Lavie

**Affiliations:** Department of Biochemistry and Molecular Genetics, University of Illinois at Chicago, Chicago, Illinois, USA; Research Resources Center, University of Illinois at Chicago, Chicago, Illinois, USA; Department of Medicinal Chemistry and Pharmacognosy, University of Illinois at Chicago, Chicago, Illinois, USA; The Jesse Brown VA Medical Center, Chicago, Illinois, USA

## Abstract

Ectonucleoside Triphosphate Diphosphohydrolase 5 (ENTPD5) has been shown to be important in maintaining cellular function in cancer, and its expression is upregulated through multiple, unique pathways in certain cancers, including laryngeal, glioblastoma multiforme, breast, testicular, and prostate. ENTPD5 supports cancer growth by promoting the import of UDP-glucose, a metabolite used for protein glycosylation and hence proper glycoprotein folding, into the ER by providing the counter molecule, UMP, to the ER antiporter. Despite its cancer-supporting function, no small molecule inhibitors of ENTPD5 are commercially available, and few studies have been performed in tissue culture to understand the effects of chemical inhibition of ENTPD5. We performed a high-throughput screen (HTS) of 21,120 compounds to identify small molecule inhibitors of ENPTD5 activity. Two hits were identified, and we performed a structure activity relationship (SAR) screen around these hits. Further validation of these probes were done in an orthogonal assay and then assayed in cell culture to assess their effect on prostate cancer cell lines. Notably, treatment with the novel ENTPD5 inhibitor reduced the amount of glycoprotein produced in treated cells, consistent with the hypothesis that ENTPD5 is important for glycoprotein folding. This work serves as an important step in designing new molecular probes for ENTPD5 as well as further probing the utility of targeting ENTPD5 to combat cancer cell proliferation.

## Introduction

Ectonucleoside Triphosphate Diphosphohydrolase 5 (ENTPD5) is the endoplasmic reticulum (ER) resident member of the NTPDase enzyme family. Unlike other members of this family, which generally catalyze the removal of the gamma and beta phosphates on triphosphate nucleotides, ENTPD5 catalyzes the removal of the terminal phosphate of UDP and GDP to form UMP and GMP, respectively[1]. This hydrolysis of UDP to UMP provides a counter molecule for the ER UDP-Glucose antiporter, which imports new UDP-glucose into the ER for proper glycoprotein folding[2].

ENTPD5 is overexpressed through two independent pathways in cancer cells. PTEN null tumors promote ENTPD5 expression via the PI3K signaling pathway, through the activation of Akt by PIP3 to p-Akt, and the sequestration of FoxO transcription family to the cytoplasm[3]. This sequestration of FoxO releases its negative regulation on ENTPD5 expression[4]. Due to the importance of ENTPD5 for the ER processing of cell surface receptors, many of which signal through the PIP3/Akt pathway, a positive feedback loop exists to accelerate ENTPD5 expression, cell growth, and glucose utilization[4] (Fig 1). The PTEN gene is at least partially deleted in 10–30% of prostate cancer tumor samples and predicts poor clinical outcomes[5–8]. ENTPD5 is also overexpressed in p53 gain-of-function mutations through interaction of Mut-p53 with Sp1 ENTPD5’s promoter region[9].

**Fig 1.**
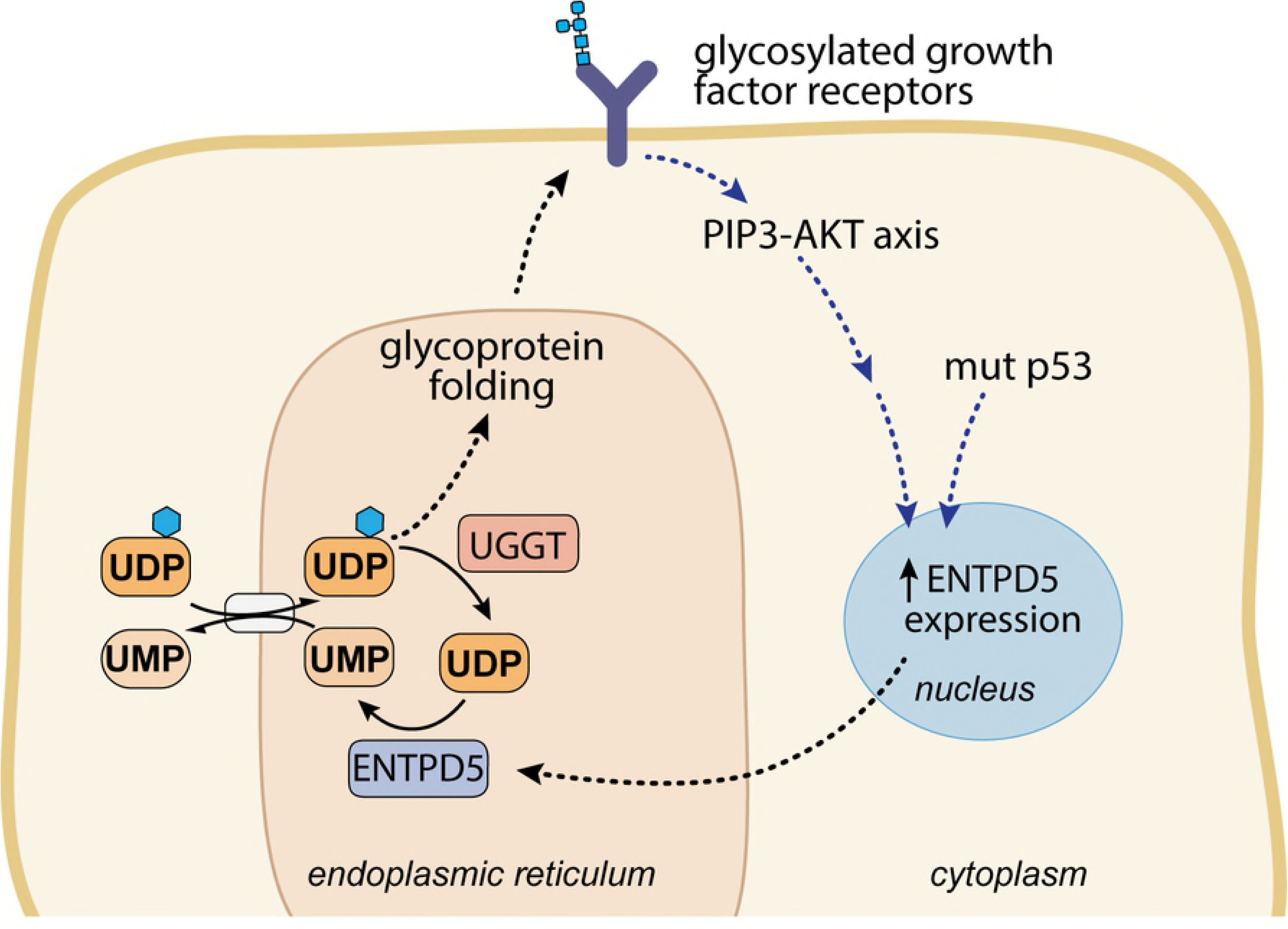
ENTPD5 is an ER-resident UDPase important for proper glycoprotein folding. Schematic diagram highlighting the role ENTPD5 plays in the glycoprotein refolding cycle in the ER. For proper glycoprotein folding to occur, UDP-glucose is brought into the ER by an antiporter that uses UMP as the counter molecule. ENTPD5 activity produces UMP, leading to increased levels of UDP-glucose entering the ER for glycoprotein refolding. ENTPD5 expression is upregulated through two independent pathways: PI3-AKT axis signaling and mutant p53 interactions.

ENTPD5 is believed to support cancer growth via two mechanisms. First, the high protein synthesis demand of cancer cells puts the protein folding machinery under stress, including the machinery to properly fold glycosylated proteins trafficked through the ER. ENTPD5 relieves ER stress in cancer cells by providing UMP for the UDP-glucose antiporter, allowing for more cycles of glycoprotein folding in the ER (Fig 1) [1, 4, 10]. Without the required post-translational glycosylation events in the ER, the unfolded protein response would be activated and the various growth factor receptors and nutrient transporters would be marked for degradation due to misfolding^[11]^. Overexpression of ENTPD5 allows cells to make large amounts of glycoproteins, including growth factor receptors and nutrient transporters that can activate a positive feedback loop through the PIP3/Akt pathway. Second, the altered metabolic state of cancer cells (i.e. the Warburg effect) affects nucleotide pools, and ENTPD5 helps to promote a balanced nucleotide pool compatible with the requirements of the cancer cell^[4]^.

ENTPD5’s promise as a target against cancer was demonstrated using inducible shRNA in xenograft models. LNCaP cells with inducible shRNA against ENTPD5 showed reduced tumor burden after knockdown[4], and MDA-MB-231 xenografts also showed reduced tumor growth following knockdown[9]. Furthermore, the viability of ENTPD5 knockout mice demonstrate the non-essentiality of this enzyme for normal tissue[10].

Currently, no small molecule inhibitors against ENTPD5 are validated in the peer-reviewed literature, though a 2010 Ph.D. thesis describes several small molecule inhibitors of ENTPD5 that show activity in cell culture at 10 μM[12]. IC_50_ values were not determined in that study or subsequent patent application. To discover additional and potentially more potent ENTPD5 inhibitors, we performed a high-throughput screen (HTS) and subsequent structure activity relationship (SAR) that identified novel small molecular probes of this important enzyme. The probes identified through HTS were further validated in an orthogonal assay and then assayed in cell culture to assess their effect on prostate cancer cell lines. These validated molecular probes provide at least two scaffolds for the further development of ENTPD5 inhibitors, and serve as an important step in probing the utility of targeting ENTPD5 to combat cancer cell proliferation.

## Materials and methods

### Materials

All chemicals were reagent or molecular biology grade. Pfu ultra polymerase (600380-51) was purchased from Agilent. 6x DNA loading dye (R0611), 1kb DNA ladder (SM1333), and 10 mM premixed dNTPs (R0192) were purchased from Thermo-Scientific. Phusion polymerase (M05305), BamHI-HF (R31365), NDE1 (R011S), and Gibson Assembly Master Mix (M5510A) were purchased from New England Biolabs. QIAquick gel extraction kit (28706) and QIAprep spin miniprep kit (27004) were manufactured by Qiagen. All oligonucleotides for PCR amplification and mutagenesis were purchased from IDT. ATP (987-65-5) was purchased from Pharma Waldhof. UDP (94330), and UMP (U6375) were purchased from Sigma-Aldrich. D-Luciferin (14681) was made by Cayman Chemical.

### Gene synthesis, cloning, protein expression, and purification of ENTPD5 from an *Escherichia coli* expression system

Codon-optimized Δ43ENTPD5 for bacterial production was synthesized by Genscript with N-terminal NdeI and C-terminal BamHI sites and cloned into a modified tag-less pET14b vector in house. Site directed mutagenesis (Forward primer: CATCTCACATGGATCCGAGAATCTTTATTTTCAGGGCCATCATCACC; Reverse primer: GGTGATGATGGCCCTGAAAATAAAGATTCTCGGATCCATGTGAGATG) using Pfu ultra was used to generate a C-terminal His_8_ tag for nickel chelate affinity purification. The resulting DNA construct (verified by Sanger sequencing) was transformed into C41(DE3) *E. coli* cells.

Cells were grown at 37 °C in 2xYT medium supplemented with 100 μg/mL ampicillin (Amp), treated with 0.1 mM isopropyl β-D-1-thiogalactopyranoside (IPTG) at an OD600 nm of 0.6–0.8, and then cultured for an additional 18h at 18 °C. Cells were harvested by centrifugation, washed with 200 mM NaCl and 25 mM Tris pH 7.5, and pelleted at 5000 rpm for 20 min before storage at -20°C.

Protein was purified from inclusion bodies using a refolding protocol modified from Murphy-Piedmonte *et al*^[13]^. After thawing, cells were lysed by sonication in 25 mM Tris pH 7.5, 200 mM NaCl, 10% glycerol, 1% Triton X-100, and 1 mM PMSF. Lysed cells were centrifuged at 20,000 rpm for 30 min. The supernatant was decanted, and the resulting pellet was resuspended in 6 M guanidine-HCl with 25 mM Tris pH 8.0 and 10 mM DTT and left to stir overnight at 4°C. The solution was centrifuged at 20,000 RPM, 4°C, and the resulting denatured protein supernatant was stored at 4°C. The denatured protein was refolded by a quick dilution method in which 25 mL of 4 mg/mL denatured protein in 6 M guanidinium solution was added to 1 L of 600 mM NaCl, 100 mM Tris-HCl pH 8.5, 2 mM reduced glutathione, 1 mM oxidized glutathione, and 10% glycerol at a rate of 0.2 mL/min at 4°C with gentle stirring for 72 hours. The protein was precipitated with 60% ammonium sulfate, centrifuged at 14,000 RPM for 1 hour, and the supernatant was discarded. The pellet was resuspended in 500 mM NaCl, 25 mM Tris pH 8.5, and 30 mM imidazole. This solution was loaded onto a 5-mL GE His-Trap column. Protein was eluted into 500 mM NaCl, 25 mM Tris-HCl pH 8.5, 250 mM imidazole. The purified protein was concentrated to 5 mL and injected onto S-200 gel filtration column (GE Healthcare) equilibrated with 250 mM NaCl, 25 mM Tris-HCl pH 8.5. To confirm the purity, collected fractions were analyzed by SDS-PAGE and detected with Coomassie Brilliant Blue staining. Activity of purified protein was assessed using the Malachite Green assay (below). All fractions containing purified active protein were pooled and stored at -80°C. The ENTPD5 purified using *E. coli* expression (referred to as B.ENTPD5) was use for the HTS.

### Gene synthesis, cloning, protein expression, and purification of ENTPD5 from baculovirus expression vector system (BEVS)/insect cells for kinetics and validation

Human cDNA for ENTPD5 with a C-terminal Flag tag was obtained from SinoBiology for BEVS and was cloned into a modified pAcGP67-A vector^[14]^. Site-directed mutagenesis was performed to introduce silent mutations in order to remove an internal EcoRI site (forward primer: AGGGAGCACTGGAACTC**GTAT**CCATGTTTACACCTTTGTG; reverse primer: CACAAAGGTGTAAACAT**GGATA**CGAGTTCCAGTGCTCCCT) and internal BamHI site (forward: GTTAGCATCATGGAT**GGCAG**CGACGAAGGCATATTAG, reverse: CTAATATGCCTTCGTC**GCTGCC**ATCCATGATGCTAAC). Initially a Δ43ENTPD5 sequence with a C-terminal His_8_ tag was amplified and inserted using the upstream BamHI site and downstream EcoRI sites, which preserved the pAcGP67-A export signal sequence in frame (forward primer: GC**GGATCC**CAGCGCCAGCACCTTGTATGG ; reverse primer: CG**GAATT**CTTAATGATGGTGATGATGGTGATGATGACCCCCATGGGAGATGCCC). The resulting DNA construct (verified by Sanger sequencing) was inserted into the Baculovirus genome using ProGreen linearized baculovirus genome kit (K20) from AB Vector according to the manufacturer. Baculuovirus stocks were amplified in SF9 cells (11496015 Gibco) cultured in Sf-900 II SFM (10902096 ThermoFisher). Six days after inoculation of 1 L of Hi-5 cells (B85502 ThermoFisher) cultured in Express Five SFM (10486025 ThermoFisher) with 100 mL of high-titer baculouvirus, the cells were centrifuged at 4,000 RPM for 25 minutes, the supernatant was filtered with a 0.22 μM filter and added to 5 mL of loose Ni-Sepharose Excel beads (17371201 GE Life Sciences). The beads were washed with 500 mM NaCl, 25 mM Tris-HCl pH 8.5, 25 mM imidazole, and the protein was eluted with 500 mM NaCl, 25 mM Tris-HCl pH 8.5, 250 mM imidazole. The purified protein was concentrated to 5 mL and injected onto a S-200 gel filtration column equilibrated with 250 mM NaCl, 25 mM Tris-HCl pH 8.5. To confirm the purity, collected fractions were analyzed by SDS-PAGE and detected with Coomassie Brilliant Blue staining. Activity of purified protein was assessed by Malachite Green assay. All fractions containing purified active protein were pooled and stored at -80 °C.

### Protein expression and purification of coupling enzymes UMP/CMPK and luciferase from *E. coli*

A UMP/CMP kinase in the pGEX-GST vector was purified as previously described[15]. Briefly, cells were grown at 37 °C in 2xYT medium supplemented with 100 μg/mL Amp. Protein expression was induced with 0.1 mM IPTG. Following overnight growth at 20 °C, cells were harvested by centrifugation, washed with 200 mM NaCl and 25 mM Tris-HCl pH 7.5, pelleted at 5,000 rpm for 20 min, and frozen. After thawing, cells were lysed by sonication in 25 mM Tris-HCl pH 8.0, 200 mM KCl, 10% glycerol, 1% Triton X-100, and 1 mM PMSF. Lysed cells were centrifuged at 20,000 rpm for 30 min. The supernatant was filtered using a 0.45 μM filter and loaded onto a 5 mL Glutathione-Sepharose column. Following a wash step with 200 mM KCl, 10 mM Tris-HCl pH 8.0, the protein was eluted with 200 mM KCl, 10 mM Tris-HCl pH 8.0, 10 mM reduced glutathione. 10 mM DTT was added to eluted protein, which was flash frozen and stored at -80°C.

Wild-type firefly luciferase in a pQE30 vector was obtained from the Branchini Lab[16]. Cells were grown at 37 °C in 2xYT medium supplemented with 100 μg/mL Amp and 25 μg/mL kanamycin. Protein expression was induced with 0.1 mM IPTG. Following overnight growth at 20 °C, cells were harvested by centrifugation, washed with 200 mM NaCl and 25 mM Tris-HCl pH 7.5, pelleted at 5,000 rpm for 20 min, and frozen. After thawing, cells were lysed by sonication in 25 mM Tris-HCl pH 7.5, 200 mM NaCl, 10% glycerol, 1% Triton X-100, and 1 mM PMSF. Lysed cells were centrifuged at 20,000 rpm for 30 min, and the supernatant was filtered through a 0.45 μM filter and loaded onto a 5-mL His-Trap column. The column was washed with 250 mM NaCl, 25 mM Tris-HCl pH 7.5, 50 mM imidazole. The protein was eluted with 250 mM NaCl, 25 mM Tris-HCl pH 7.5, 250 mM imidazole. Eluted protein was dialyzed into 150 mM NaCl, 50 mM Tris pH 7.5, 1 mM DTT, and 1 mM EDTA, flash frozen, and stored at -80°C.

### High throughput Screening

Stocks of purified B.ENTPD5, UMPK, and luciferase were diluted to 0.4 mg/mL, 1.2 mg/mL, 4.5 mg/mL, respectively, in 250 mM NaCl, 25 mM Tris-HCl pH 8.5, 5 mM MgCl_2_, 0.1 mg/mL BSA, and 5 mM DTT and flash frozen. To prepare enzyme mix solution for HTS, ENTPD5 (1 ng/μL final concentration) and UMPK (3.5 ng/μL) were diluted in HTS buffer (250 mM NaCl, 25 mM Tris-HCl pH 7.7, 5 mM MgCl_2_, 5 mM DTT, 0.1 mg/mL BSA, and 0.01% Triton X-100). An ENTPD5 null mix was made as above without ENTPD5 to provide maximal signal mimicking 100% inhibition. Substrate mix solution consisted of ATP (625 μM) and UDP (150 μM) in HTS buffer. Developer solution consisted of Luciferase (50 ng/μL) and D-luciferin (1.6 mM) in HTS buffer. All solutions were stored at 4°C, and equilibrated to room temperature prior to screening.

All steps of ENTPD5 HTS were performed at room temperature in duplicate 384-well plates. For each run, 40 μL of enzyme solution (or ENTPD5 null mix) was added to each well of a 384 well plate. 0.1 μL of 10 mM compound (in 100% DMSO) from a 25,000 subset of the Chembridge DiverSet library or DMSO alone were added to wells by pin tool (V&P Scientific). Following a 10-minute incubation, the reaction was initiated with 10 μL of substrate mix (final concentrations: 40 ng/50 μL ENTPD5, 140 ng/50 μL UMPK, 125 μM ATP, 30 μM UDP, 20 μM test compound). Assay plates were shaken at 1500 rpm for 30 seconds and then incubated for 1h. Following incubation, 10 μL of developer solution was added to each well (final concentrations: 500 ng/60 μL luciferase, 266 μM D-luciferin). The plate was shaken for 30 seconds and incubated for 30 minutes. Luminescence readings were performed on a Tecan Infinite F200Pro with automatic attenuation settings and 100 ms integration time.

Hits and hit analogs were (re)purchased from Chembridge for SAR studies. Compounds were assayed at 5 μM, 10 μM, 20 μM, and 40 μM using the assay described above.

### Malachite green assay

Malachite Green (MG) assay[17] for free phosphate was used to directly test ENTPD5 activity. Standard curves for each preparation of malachite green were generated using known amounts of free phosphate. Reactions were carried out at room temperature in 1-mL cuvettes and OD 630 nm measurements were performed on a Nanodrop 2000c. ENTPD5 assays were initiated by adding UDP (100 μM final concentration) to a cuvette containing 800 μL of ENTPD5 (10 ng) in reaction buffer (250 mM NaCl, 5 mM MgCl_2_, 25 mM Tris-HCl pH 8.5). After 5 minutes, the reaction was quenched with 200 μL of MG buffer (13.1% (v/v) Concentrated sulfuric acid, 2.64 mM malachite green, 1.4% (w/v) ammonium molybdate, and 0.17% (v/v) Tween 20) and mixed by pipet. The OD 630 nm was measured after an additional 5-minute incubation. IC_50_ values were calculated by GraphPad Prism 6 using sigmoidal interpolation model with 95% confidence intervals.

### Cell culture

LNCaP (ATCC CFL-1740) and DU-145 (ATCC HTB-81) cell lines were cultivated in a humid atmosphere (5% CO2, 37°C) using RPMI 1640 with L-glutamine (10-040-CV Corning) media supplemented with 10% FBS (SH30910.03 Hyclone) and 1x penicillin-streptomycin solution (Invitrogen). All cell lines were analyzed by STR (Short Tandem Repeat) and confirmed to match to corresponding STR profile data from the Global Bioresource Center ATCC or ExPASy Cellosaurus database. All cell lines were verified to be mycoplasma free.

### Growth inhibition assay

5,000 cells were plated on day 0 in a 96-well plate. On day 1, before treatment, each well was imaged and direct cell counting was performed using a Celigo S (Nexcelom) imager and automatic cell counting software provided with the instrument. Compounds were added to final concentrations of 0.15 μM- 20 μM. 48 hours after treatment, wells were re-imaged and counted using the pre-treatment counting procedure. Growth was reported as the fold increase over the baseline measurement and normalized to DMSO controls. Experiments were performed in triplicate, in three separate experiments. EC_50_ values were calculated by GraphPad Prism 6 using a sigmoidal interpolation model with 95% confidence intervals.

### Western Blots

LNCaP and DU145 cells were treated with 10 μM compound or DMSO control. 24h post-treatment, cells were washed with ice cold PBS, scraped, pelleted and stored at -80°C until use. Frozen cell pellets were lysed with lysis buffer consisting of 20 mM HEPES pH. 7.4, 150 mM NaCl, 1% Triton X-100, 1 mM EGTA, 1 mM EDTA, 10 mM sodium pyrophosphate, 100 mM NaF, 5 mM iodoacetic acid, 20 nM okadaic acid, 0.2 mM phenylmethylsulfonyl flouride (PMSF), and complete protease inhibitor cocktail tablets (Roche Diagnostics). Samples containing 50 μg of protein were suspended in SDS loading buffer, separated on 4–12% SDS polyacrylamide gels (GenScript #M41215) and electrotransferred to PVDF membranes (Millipore Immobilon-FL #IPFL00010). Precision Plus Protein Standards all blue (Bio-Rad 161-0373) were included as molecular weight markers. Immunoblotting was performed using standard methods, with TBS-T and TBS-T with 5% BSA as the wash and blocking/primary antibody dilution solutions, respectively. Total protein was measured with Ponceau-S staining prior to probing with antibodies. Membranes were incubated overnight at 4°C with primary antibodies diluted 1:1000 unless noted otherwise. Antibodies included: mouse monoclonal anti-ENTPD5 (Santa Cruz Biotechnology – sc-377172, 1:100 dilution), rabbit monoclonal anti-PTEN (Cell Signaling - 138G6), rabbit monoclonal anti-Sp1 (abcam - ab124804), rabbit monoclonal anti-GAPDH (Cell Signaling - 2118), and mouse monoclonal anti-O-GlcNAc (BioLegend - 838004). Membranes were incubated with 1:20,000 dilutions of IRDye 800CW Goat anti-Mouse IgG (LI-COR #925-32210) and IRDye 680RD Goat anti-Rabbit IgG (LI-COR #925-68071) for 1 h at room temperature, visualized on a LI-COR CLx near IR scanner. Western blot quantification was normalized to total protein levels and analyzed in ImageStudio (LI-COR).

## Results

### Recombinant expression and purification of ENTPD5

An N-terminal truncation construct of ENTPD5 was recombinantly expressed in *E. coli* as well as in insect cells using the baculovirus expression vector system (BEVS). The protein was expressed as a 43-residue N-terminal truncation to more closely mimic the mature protein resulting from cleavage of its ER signal sequence (residues 1–24). The deletion was extended to residue 43 after analysis of sequence alignments with other members of the ENTPD family across species as well as the known crystal structures of rat ENTPD1[18] and ENTPD2[19] to avoid potential complications with a cysteine residue at position 39. The structure of ENTPD5 is predicted to contain two conserved disulfide bonds (Fig 2a). Cysteine-39 is a non-conserved cysteine that is predicted to not be required for proper protein folding, and its absence should reduce the propensity for the formation of incorrect disulfide bridges, especially in the process of refolding the enzyme. For bacterial expression, bacterial codon-optimized human ENTPD5 44-428 (B.ENTPD5) was cloned into pET14b vector with a C-term His8-tag (Fig 2a). Recombinant ENTPD5 was also produced in insect cells (I.ENTPD5) using BEVS. This expression system produced natively active protein without denaturing, refolding, or shuffling of disulfide bonds.

**Figure 2:**
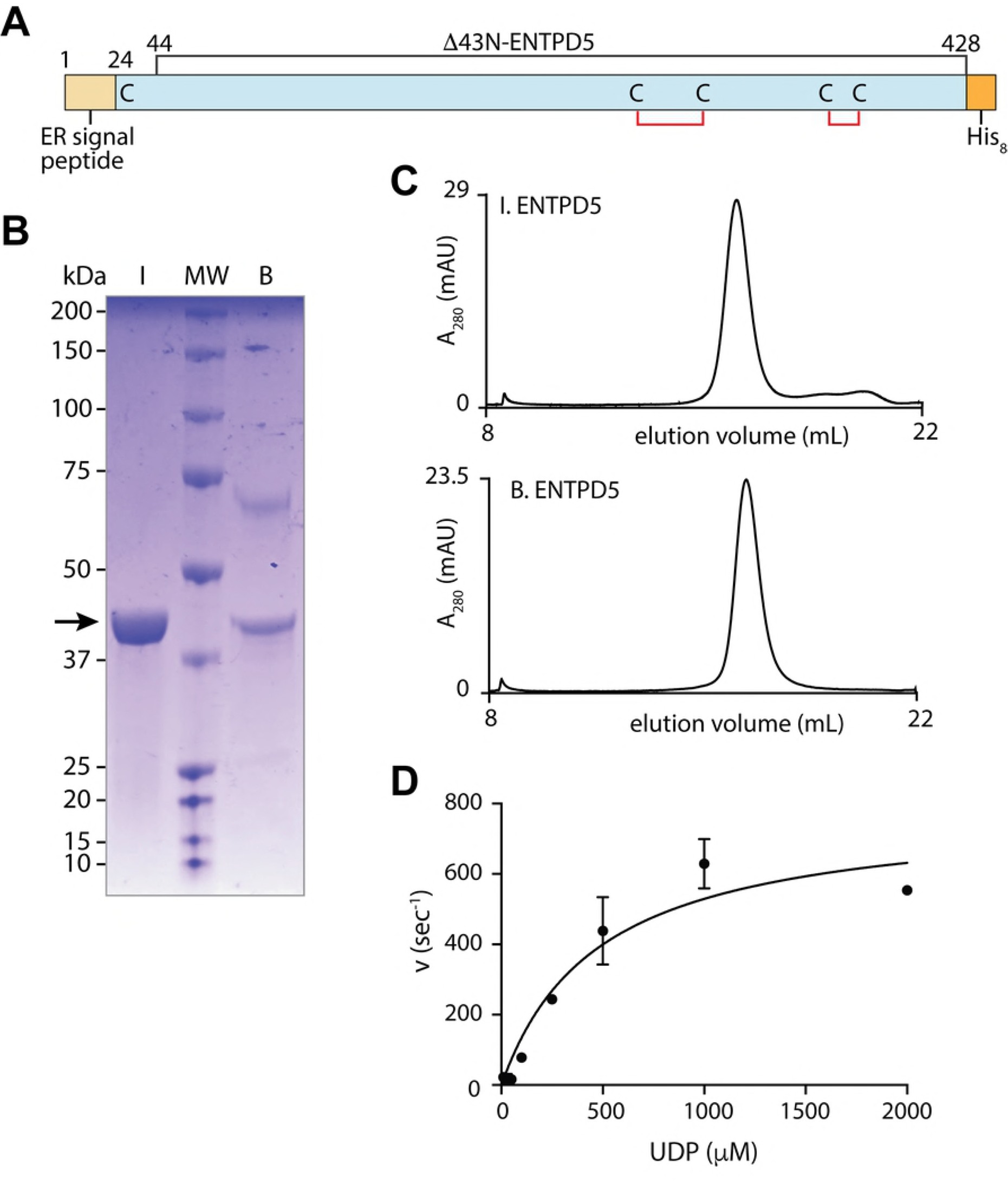
ENTPD5 construct, purification, and activity. **A**) Diagram of ENTPD5 construct used for recombinant protein production, highlighting the expected internal disulfide bonds. **B**) SDS-PAGE gel of ENTPD5 purified from insect cells (lane **I**) and from bacteria (lane **B**). Molecular weight marker (lane **MW**) sizes in kDa are shown to the left of the gel. BSA, the additional band in lane **B** at 66 kDa, was added post-purification to stabilize B.ENTPD5. **C**) Gel filtration traces of recombinant ENTPD5 proteins. **D**) Michaelis-Menten plot of purified I.ENTPD5 activity with substrate UDP using the Malachite Green assay. K_m_: 480 ± 111 μM; *k*_cat_: 783 ± 68 s^−1^. Reactions were performed in triplicate. Error bars designate standard deviation.

Both sources of purified protein appear the same size on SDS-PAGE (Fig 2b) and behave similarly on gel filtration, corresponding to the expected monomeric size (Fig 2c). The ~65 kDa band observed in the B.ENTPD5 sample is BSA added post-purification for protein stability purposes. Maximum enzymatic activity was achieved using the I.ENTPD5 protein (Fig 2d), corresponding to a ~30% increase in *k*_cat_ relative to that previously reported for refolded hENTPD5 purified from bacteria^[13]^.

### High-throughput screening to identify inhibitors of ENTPD5

To discover ENTPD5 inhibitors, a coupled enzyme assay was developed for high-throughput screening. The ENTPD5 reaction (UDP → UMP + Pi) was coupled with uridine monophosphate kinase’s (UMPK) reaction with UMP (UMP + ATP → UDP + ADP) to create a futile catalytic cycle consuming ATP (Fig 3a). After one hour, the residual ATP was measured with firefly luciferase. UMPK concentrations were chosen to ensure ENTPD5, not UMPK, was rate-limiting. The ATP concentration was chosen to maximize signal with no ENTPD5 present, and the reaction time was optimized to remain in the linear range of the assay. Initial UDP concentration was chosen to allow for multiple turnover events on a single molecule in the cycle. Due to the availability of higher yields of the bacterially produced enzyme, B.ENTPD5 was used for HTS and subsequent structure activity relationship studies. 21,000 compounds were screened in duplicate at 20 µM. The average Z’ value of the 132-plate screen was 0.73, indicating a robust assay[20]. Five compounds producing >25% reproducible inhibition of ENTPD5 activity were selected for further analysis (0.02% hit rate) (Fig 3b, c).

**Figure 3:**
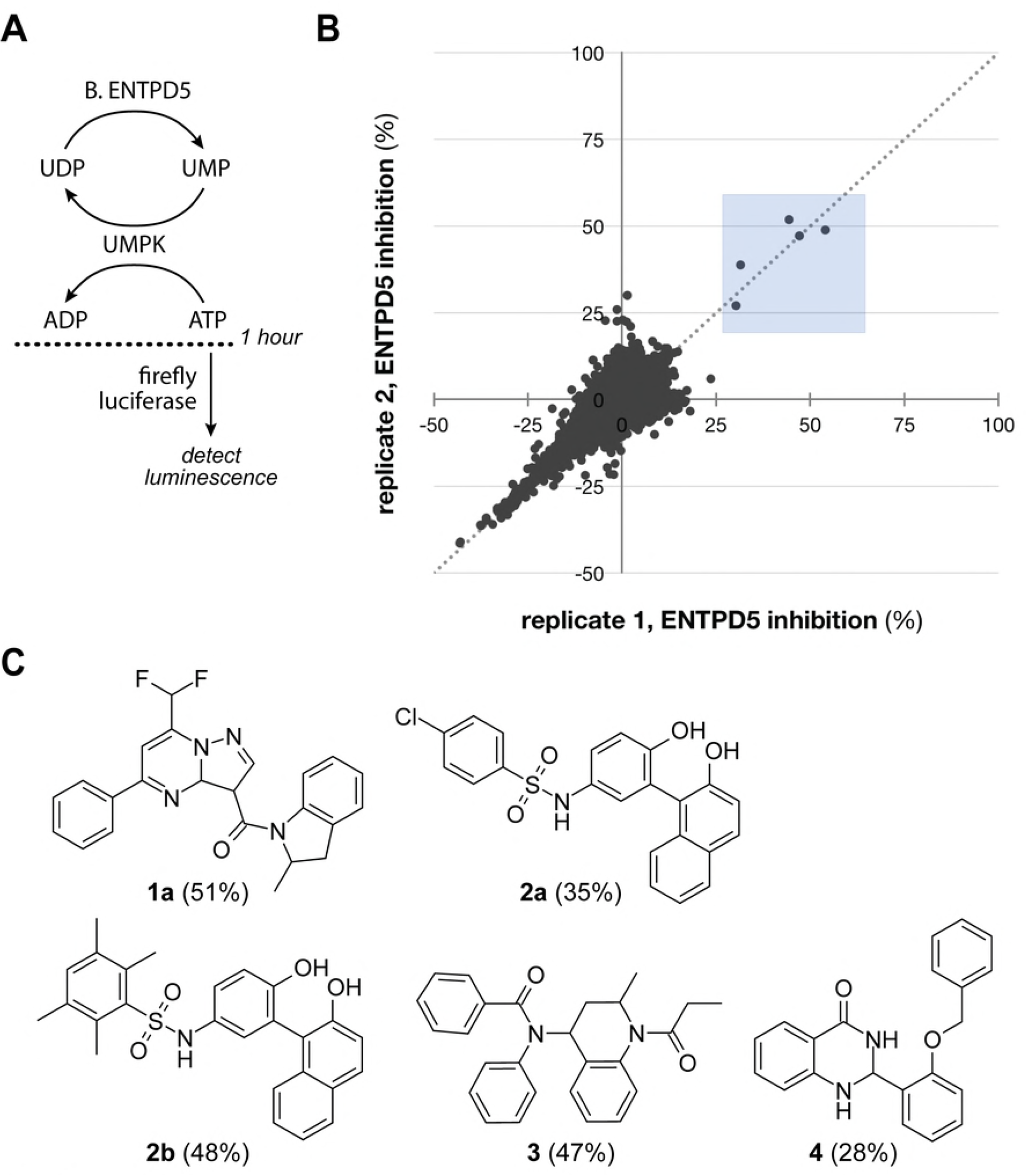
ENTPD5 HTS assay and screening results. **A**) Schematic of ENTPD5 HTS assay. Following a 1-hour coupled reaction of ENTPD5 and UMPK, the residual ATP is measured indirectly using luciferase. **B**) Replicate plot of ENTPD5 HTS, showing percent ENTPD5 inhibition by compounds in each replicate set. Compounds were screened at 20 µM in duplicate. Compounds producing >25% inhibition (blue shaded box in **B**) were selected for follow-up analysis and are shown in **C**). Percent ENTPD5 inhibition by each hit is shown in parentheses.

### Validation of ENTPD5 HTS hits

Using I.ENTPD5, the five hits compounds were validated in the malachite green (MG) assay[17], which directly measures free phosphate, a product of ENTPD5 activity. Compounds **1a**, **2a**, **2b**, and **4** displayed a dose-dependent inhibition of I.ENTPD5.

Hit compounds **1a**, **2a**, and **2b** were used as the basis for structure activity relationship studies, in which ten analogs that were at least 80% structurally similar to the initial hits were tested against ENTPD5 at 5, 10, 20 and 40 µM (Fig 4). Analogs of compound **4** were not available from the supplier at the time of SAR. All three analogs of compound **1a** were inactive against ENTPD5, highlighting the importance of a methyl group at position R^2^. The compound **2** series SAR demonstrates that substitutions on the benzenesulfonamide are well tolerated (Fig 4, core A series). However, replacing the N-hydroxyphenyl group with an N-hydroxynapthalene leads to a drastic loss in potency (Fig 4, core B series).

**Fig 4.**
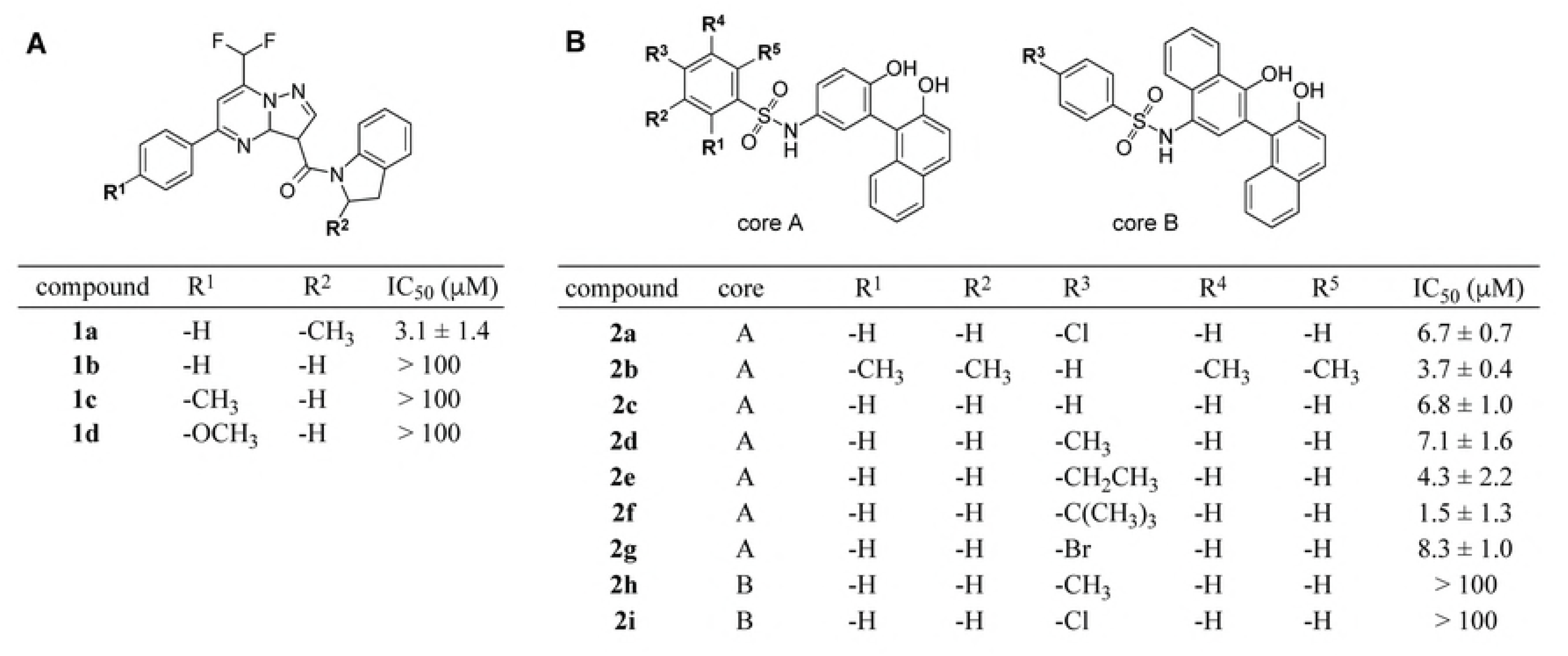
Structure activity relationship of select ENTPD5 inhibitors identified by HTS. Close analogs of hits **1a** (A) and **2a** (B) were assayed with B.ENTPD5 to assess the effect of conservative substitutions to the scaffolds. Compounds were assayed at 5, 10, 20, and 40 μM using the coupled enzyme HTS assay to determine IC_50_ values. Data were fit by nonlinear regression.

The most potent compounds from each inhibitor scaffold, **1a** and **2f**, were re-assayed in the MG assay at an expanded concentration range. Limited solubility of the hits prevented us from achieving a concentration required for full inhibition of the ENPTD5 activity. Nevertheless, these studies revealed IC_50_ values of 3.14 µM and 1.54 µM for compounds **1a** and compound **2f**, respectively, and IC_50_ >100 µM for inactive compound **1b** (Fig 5).

**Figure 5:**
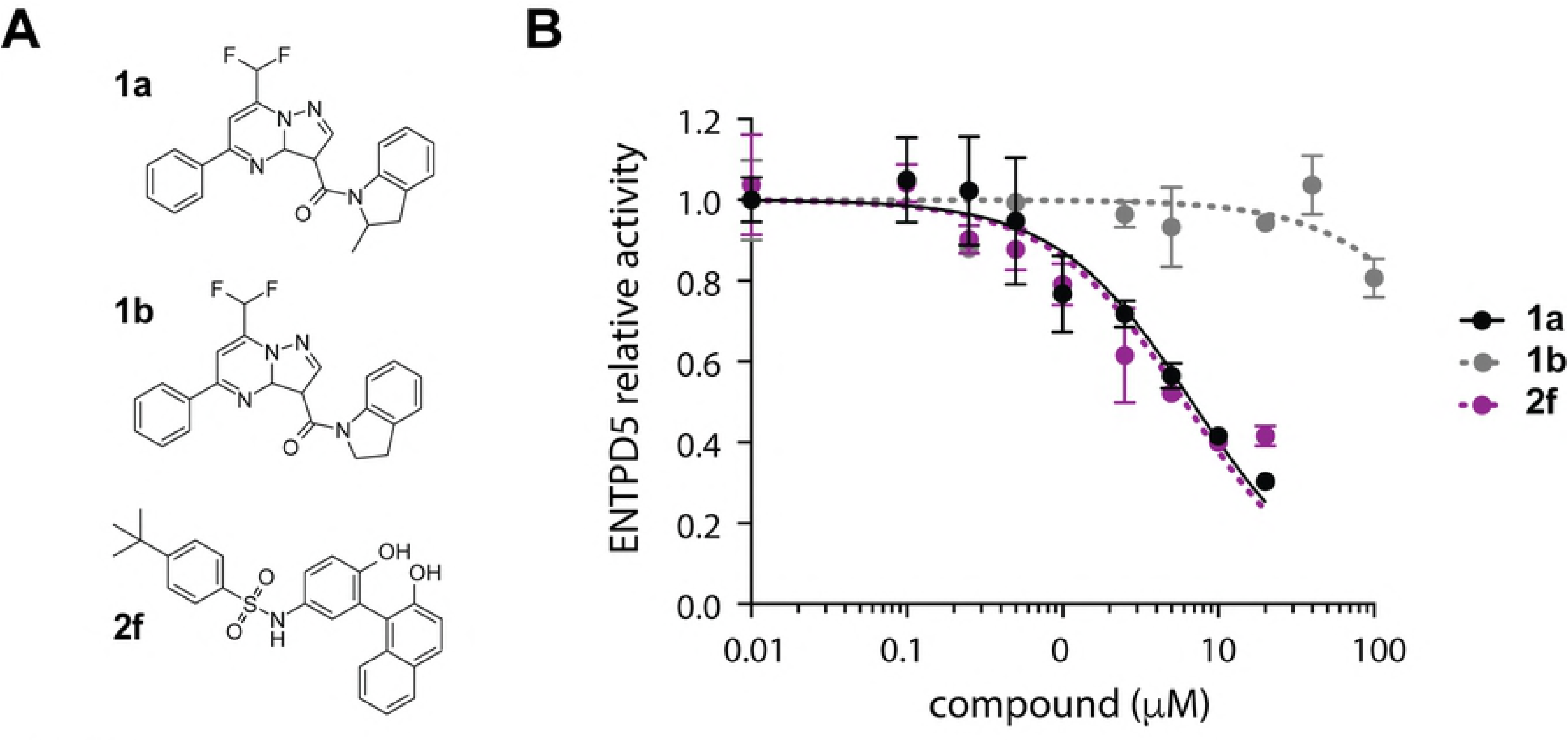
ENTPD5 HTS hit characterization. **A)** Structures of compounds **1a, 1b**, and **2f**. **B)** ENTPD5 inhibition by compounds **1a, 1b**, and **2f** using the MG assay. **1a** IC_50_: 3.1 ± 1.4 μM; **1b** EC_50_ >100 μM; **2f** EC_50_: 1.5 ± 1.3 μM. Error bars represent standard deviation from triplicate measurements.

### Inhibition of prostate cancer cell growth by compound 1

Since previous work related ENTPD5 levels to prostate cancer survival[4, 21], compounds **1a**, **1b** (as control), and **2f** were assayed with prostate cancer cell line LNCaP to determine their effect on cell proliferation. Treatment of LNCaP cells with **1a** for 48h drastically reduced the cell count, producing an EC_50_ of 0.47 µM (Fig 6a). As expected, inactive analog **1b** was much less potent against LNCaP cells (EC_50_ > 10 µM), however we were surprised to see that compound **2f** also was not effective in cell culture, with an EC_50_ > 10 uM. Compound **1a** was also assayed in an additional prostate cancer line, DU145, to investigate the effect of ENTPD5 inhibition on cell lines with differing modes of ENTPD5 overexpression. LNCaP cells are PTEN null, while both DU145 and LNCaP cells have high levels of Sp1 transcription factor[22]. After 48 hours of treatment, both cell lines were affected by inhibitor treatment. EC_50_ values of 0.47 µM and 3.12 µM were obtained for LNCaP and DU145 cells, respectively (Fig 6b).

**Figure 6:**
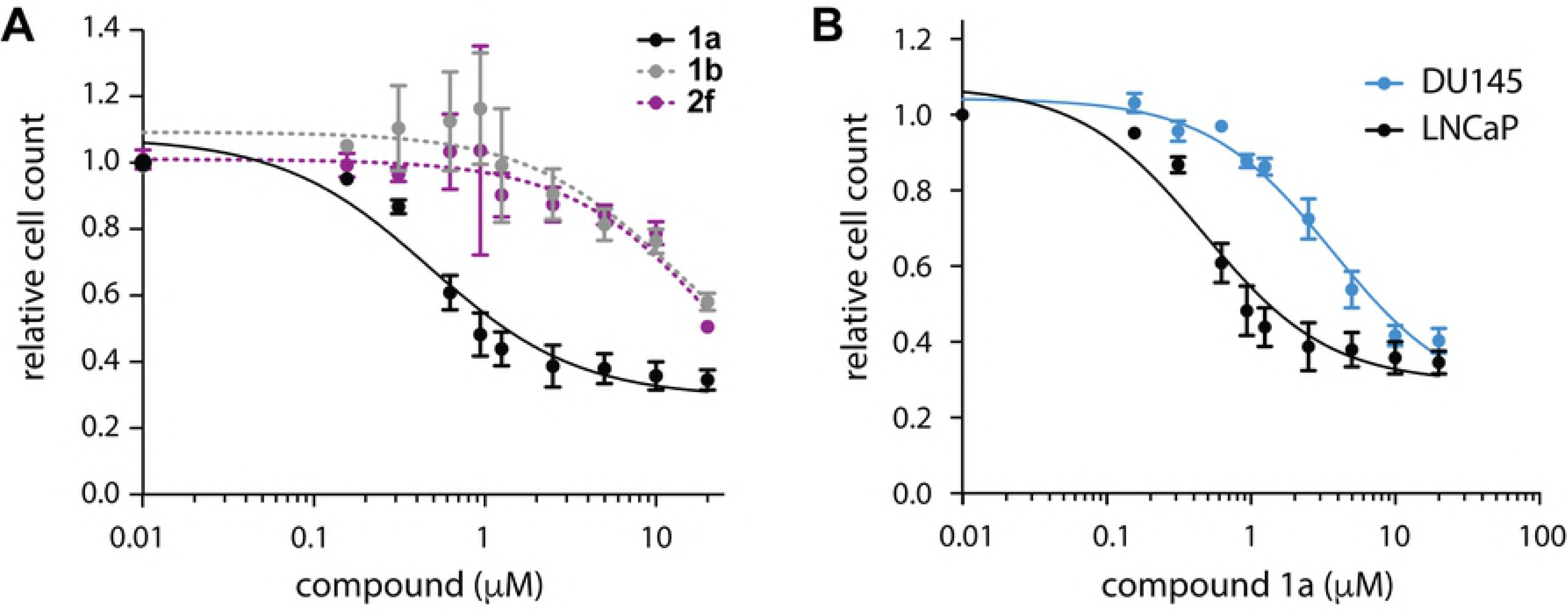
Treatment of prostate cancer cell lines with ENTPD5 inhibitors. **A)** Growth of LNCaP cells treated with compound **1a**, **1b**, or **2f** for 48h. **1a** EC_50_: 0.47 ± 1.28 μM; **1b** EC_50_: 12.9 ± 3.3 μM; **2f** EC_50_: 24 ± 3 μM. **B)** Growth of LNCaP and DU145 cells treated with compound **1a for 48h**. **1a** EC_50_ in DU145: 3.6 ± 1.2 μM; **1a** EC_50_ in LNCaP: 0.47 ± 1.28 μM. Relative cell count was normalized to DMSO-treated cell counts. Error bars represent standard error of the mean between three independent experiments.

To test the hypothesis that inhibition of ENTPD5 has an impact on cell growth due to the reduction of protein glycosylation and to further investigate the utility of **1a** as an effective molecular probe, we immunoblotted for total O-glycosylation levels within LNCaP and DU145 cells following 24 hours of treatment at 10 µM of compound **1a** and compound **2f**. First we verified that untreated LNCaP and DU145 cells have high levels of ENTPD5, Sp1, and O-glycosylation (Fig 7a). Compound **1a**, but not compound **2f**, reduced the amount of O-glycan present in the two prostate cell lines (Fig 7b, c). The decrease in O-glycan levels by compound **1a** is consistent with the effects of ENTPD5 inhibition on the glycoprotein refolding cycle. The inability of compound **2f** to reduce O-glycan levels could be attributed to low cell permeability but requires further investigation.

**Figure 7:**
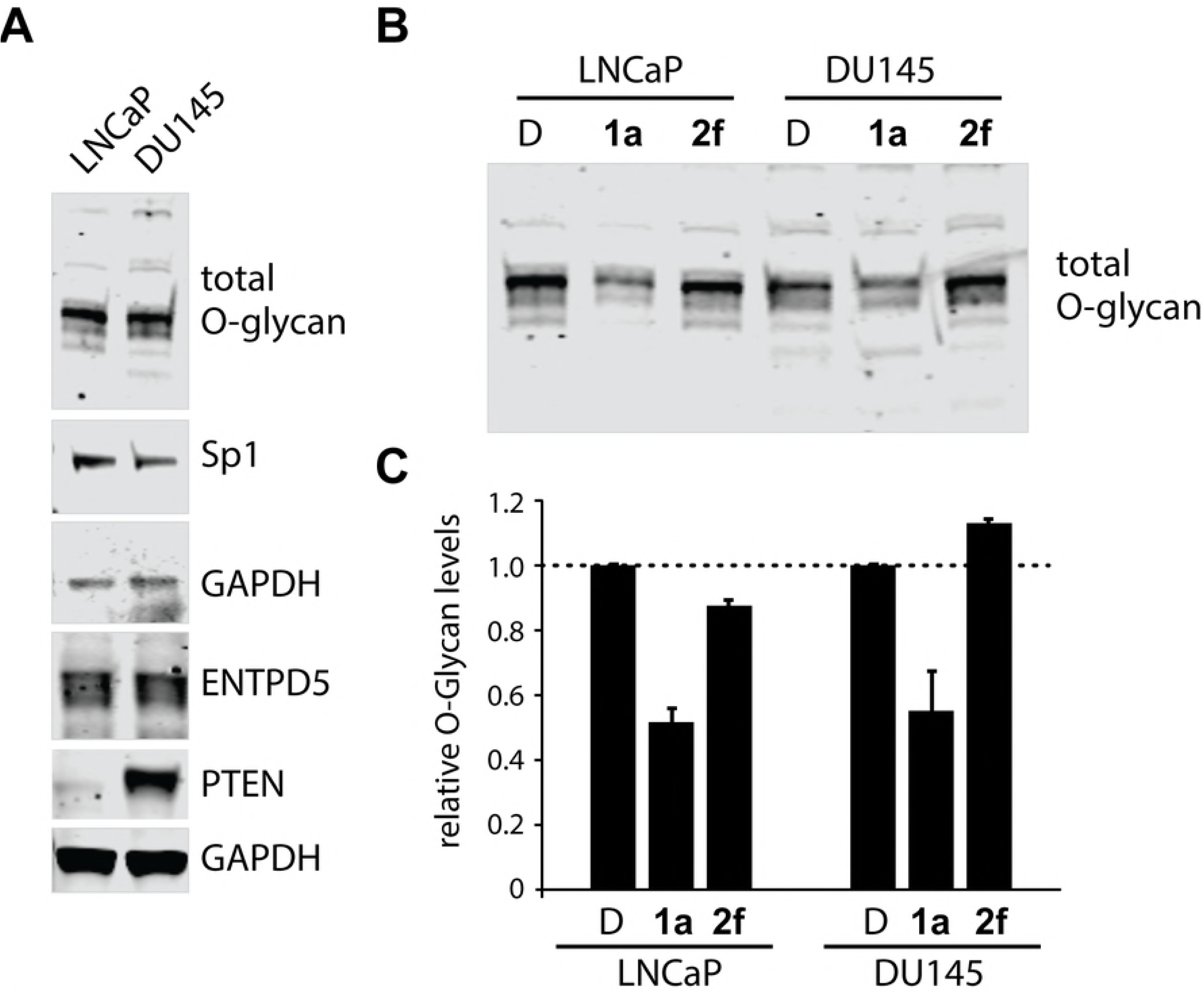
O-glycan levels of cells treated with ENTPD5 inhibitors. **A)** Baseline levels of ENTPD5, PTEN, Sp1, and O-glycans in untreated LNCaP and DU145 cells. **B)** Levels of O-glycan in LNCaP and DU145 cells treated with 10 µM **1a** or **2f** for 24h. Relative protein amounts were normalized to DMSO-treated cells from each cell line. Error bars represent standard deviation between 2 experiments.

## Discussion

Many of the newer anti-cancer approaches target a cancer driver. That is, the drugs target the oncogene that “drives” cancer growth, either as a growth factor receptor or a part of a downstream signaling cascade. However, these therapies are plagued by drug resistance, often due to the emergence of an alternative cancer driver or signaling cascade[23]. We postulate that targeting a factor that is required for cancer growth, irrespective of the cancer driver, would circumvent this type of resistance due to the natural tumor heterogeneity. In other words, as an alternative for targeting a cancer driver, which can be circumvented by other drivers in the heterogeneous cancer cell pool, we suggest targeting a cancer phenotype that is present in cancer cells regardless of any specific driver.

Indeed, several such cancer phenotypes exist, but some make for better targets than others. For example, the increased need for DNA metabolites to allow for the accelerated proliferation rate is a cancer phenotype. This phenotype is the basis for many current anti-cancer therapeutics, such as methotrexate and 5-fluoruracil, that target DNA replication. However, since many normal cells also have a high need for DNA replication, this class of drugs is associated with high toxicity. The challenge then is to identify a cancer phenotype that is largely absent in normal cells.

The fact that cancer cells have different metabolic needs outside of DNA replication generates a cancer phenotype that potentially can be targeted selectively. One component of this altered metabolism is increased expression of growth factor receptors and nutrient transporters. These membrane proteins undergo post-translational glycosylation in the ER before trafficking to the plasma membrane. In prostate cancer cells, the enzyme ENTPD5 plays a key role in maintaining the metabolite pool required for glycosylation. Importantly, ENTPD5 is not a prostate cancer driver, but rather ENTPD5 has been shown to be required for supporting prostate cancer growth. The important role ENTPD5 plays in prostate cancer proliferation is consistent with this enzyme being highly expressed in prostate cancer compared to normal prostate epithelium. Thus, an inhibitor of ENTPD5 would provide an innovative strategy to target the metabolic machinery that supports cancer growth instead of targeting a prostate cancer driver.

This paper lays the groundwork for future development of specific ENTPD5 inhibitors and molecular probes. The coupled HTS strategy that we developed was able to identify small molecules that were validated in a secondary enzyme activity assay and that show promising anti-proliferative activity against two prostate cancer cell lines. This highlights the potential of this strategy to be used for expanding HTS of ENTPD5 to larger compound collections. To our knowledge, the inhibitors identified here are the first direct small molecule inhibitors of ENTPD5 in the peer-reviewed literature. Further studies with these and additional ENTPD5 inhibitors will help to clarify whether ENPTD5 is a suitable new target for cancer therapy.

## Supporting Information

**S1 Fig. 1. Uncropped blots and ponceau stains of western blot images**. **A)** Ponceau for Fig. 6A ENTPD5 & PTEN. **B)** Uncropped image for Fig. 6A ENTPD5 on the 800 nm channel. **C)** Uncropped image for 6A PTEN (upper box) and GAPDH (lower box) in the 700 nm. **D)** Ponceau for Fig. 6A O-Glycan. **E)** Uncropped image for Fig. 6A O-Glycan on the 800 nm channel. **F)** Uncropped image for Fig. 6A SP1 (upper box) and GAPDH (lower box) on the 700 nm channel. **G)** Ponceau for Fig. 6B. **H)** Uncropped image for Fig. 6B O-Glycan on the 800 nm channel. Boxes represent area cropped for figures in the manuscript.

## References

1. Trombetta ES, Helenius A. Glycoprotein reglucosylation and nucleotide sugar utilization in the secretory pathway: identification of a nucleoside diphosphatase in the endoplasmic reticulum. EMBO J. 1999 Jun 15;18(12):3282–92. PubMed PMID: 10369669. Pubmed Central PMCID: 1171409.

2. Hirschberg CB, Robbins PW, Abeijon C. Transporters of nucleotide sugars, ATP, and nucleotide sulfate in the endoplasmic reticulum and Golgi apparatus. Annu Rev Biochem. 1998;67:49–69. PubMed PMID: 9759482.

3. Tzivion G, Dobson M, Ramakrishnan G. FoxO transcription factors; Regulation by AKT and 14–3–3 proteins. Biochim Biophys Acta. 2011 Nov;1813(11):1938–45. PubMed PMID: 21708191.

4. Fang M, Shen Z, Huang S, Zhao L, Chen S, Mak TW, et al. The ER UDPase ENTPD5 promotes protein N-glycosylation, the Warburg effect, and proliferation in the PTEN pathway. cell. 2010 Nov 24;143(5):711–24. PubMed PMID: 21074248.

5. Vidotto T, Tiezzi DG, Squire JA. Distinct subtypes of genomic PTEN deletion size influence the landscape of aneuploidy and outcome in prostate cancer. Mol Cytogenet. 2018;11:1. PubMed PMID: 29308088. Pubmed Central PMCID: 5753467.

6. Picanco-Albuquerque CG, Morais CL, Carvalho FL, Peskoe SB, Hicks JL, Ludkovski O, et al. In prostate cancer needle biopsies, detections of PTEN loss by fluorescence in situ hybridization (FISH) and by immunohistochemistry (IHC) are concordant and show consistent association with upgrading. Virchows Arch. 2016 May;468(5):607–17. PubMed PMID: 26861919.

7. Phin S, Moore MW, Cotter PD. Genomic Rearrangements of PTEN in Prostate Cancer. Front Oncol. 2013;3:240. PubMed PMID: 24062990. Pubmed Central PMCID: 3775430.

8. Verhagen PC, van Duijn PW, Hermans KG, Looijenga LH, van Gurp RJ, Stoop H, et al. The PTEN gene in locally progressive prostate cancer is preferentially inactivated by bi-allelic gene deletion. J Pathol. 2006 Apr;208(5):699–707. PubMed PMID: 16402365.

9. Vogiatzi F, Brandt DT, Schneikert J, Fuchs J, Grikscheit K, Wanzel M, et al. Mutant p53 promotes tumor progression and metastasis by the endoplasmic reticulum UDPase ENTPD5. Proc Natl Acad Sci U S A. 2016 Dec 27;113(52):E8433–E42. PubMed PMID: 27956623. Pubmed Central PMCID: 5206569.

10. Read R, Hansen G, Kramer J, Finch R, Li L, Vogel P. Ectonucleoside triphosphate diphosphohydrolase type 5 (Entpd5)-deficient mice develop progressive hepatopathy, hepatocellular tumors, and spermatogenic arrest. Vet Pathol. 2009 May;46(3):491–504. PubMed PMID: 19176496.

11. Hetz C, Chevet E, Harding HP. Targeting the unfolded protein response in disease. Nat Rev Drug Discov. 2013 Sep;12(9):703–19. PubMed PMID: 23989796.

12. Huang S. Small Molecule Regulator of ENTPD5, and ER Enzyme in the PTEN/AKT Pathway [Doctoral]: University of Texas Southwestern Medical Center at Dallas; 2010.

13. Murphy-Piedmonte DM, Crawford PA, Kirley TL. Bacterial expression, folding, purification and characterization of soluble NTPDase5 (CD39L4) ecto-nucleotidase. Biochim Biophys Acta. 2005 Mar 14;1747(2):251–9. PubMed PMID: 15698960.

14. Antanasijevic A, Kingsley C, Basu A, Bowlin TL, Rong L, Caffrey M. Application of virus-like particles (VLP) to NMR characterization of viral membrane protein interactions. J Biomol NMR. 2016 Mar;64(3):255–65. PubMed PMID: 26921030. Pubmed Central PMCID: 4826305.

15. Segura-Pena D, Sekulic N, Ort S, Konrad M, Lavie A. Substrate-induced conformational changes in human UMP/CMP kinase. J Biol Chem. 2004 Aug 6;279(32):33882–9. PubMed PMID: 15163660.

16. Sundlov JA, Fontaine DM, Southworth TL, Branchini BR, Gulick AM. Crystal structure of firefly luciferase in a second catalytic conformation supports a domain alternation mechanism. Biochemistry. 2012 Aug 21;51(33):6493–5. PubMed PMID: 22852753. Pubmed Central PMCID: 3425952.

17. Baykov AA, Evtushenko OA, Avaeva SM. A malachite green procedure for orthophosphate determination and its use in alkaline phosphatase-based enzyme immunoassay. Analytical Biochemistry. 1988;171(2):266–70.

18. Zebisch M, Krauss M, Schafer P, Strater N. Crystallographic evidence for a domain motion in rat nucleoside triphosphate diphosphohydrolase (NTPDase) 1. J Mol Biol. 2012 Jan 13;415(2):288–306. PubMed PMID: 22100451.

19. Zebisch M, Baqi Y, Schafer P, Muller CE, Strater N. Crystal structure of NTPDase2 in complex with the sulfoanthraquinone inhibitor PSB-071. J Struct Biol. 2014 Mar;185(3):336–41. PubMed PMID: 24462745.

20. Zhang JH, Chung TDY, Oldenburg KR. A simple statistical parameter for use in evaluation and validation of high throughput screening assays. Journal of Biomolecular Screening. 1999 Apr;4(2):67–73. PubMed PMID: WOS:000080223000005. English.

21. Horak P, Tomasich E, Vanhara P, Kratochvilova K, Anees M, Marhold M, et al. TUSC3 loss alters the ER stress response and accelerates prostate cancer growth in vivo. Sci Rep. 2014;4:3739. PubMed PMID: 24435307. Pubmed Central PMCID: 3894551.

22. Hay CW, Hunter I, MacKenzie A, McEwan IJ. An Sp1 Modulated Regulatory Region Unique to Higher Primates Regulates Human Androgen Receptor Promoter Activity in Prostate Cancer Cells. PLoS One. 2015;10(10):e0139990. PubMed PMID: 26448047. Pubmed Central PMCID: 4598089.

23. Neel DS, Bivona TG. Resistance is futile: overcoming resistance to targeted therapies in lung adenocarcinoma. NPJ Precis Oncol. 2017;1. PubMed PMID: 29152593. Pubmed Central PMCID: 5687582.

